# Drug-induced cytotoxicity prediction in muscle cells, an application of the Cell Painting assay

**DOI:** 10.1101/2024.02.08.579439

**Authors:** Roman Lambert, Pablo Aparicio, Eva Serrano Candelas, Aisling Murphy, Rafael Gozalbes, Howard Fearnhead

## Abstract

*In silico* toxicity prediction offers the chance of reducing or replacing most animal testing through the integration of large experimental assay datasets with the appropriate computational approaches. The use of Cell Painting to detect various phenotypic changes induced by chemicals is emerging as a powerful technique in toxicity prediction. However, most Cell Painting approaches use cancer cells that are less relevant for many toxicological endpoints, which may limit the usefulness of this data. In this study, a myoblast cell line is used to characterize cellular responses to a panel of 30 known myotoxicants. In place of traditional structural descriptors, here each perturbation is described by a fingerprint of calculated properties, deducted from the intensity, shape, or texture of individual cells. We show that these kinds of descriptors convey information to allow the prediction of the cellular viability and fate of cells in myoblasts and differentiated myotubes of the C2C12 cell line, and the clustering of drugs by their cytotoxicity responses.

**Author Summary:** Studying the toxicity of chemical compounds and drugs is crucial to avoid potentially lethal adverse effects of commercialized products, but also to detect the unsuspected toxicity of existing drugs. While these assays traditionally rely on animal models raising important ethical concerns, a need for *in vitro* and *in silico* models is present and increasing in recent years. We here propose a predictive model capable of predicting the values of a cell viability assay using cell morphology profiles captured with a microscopy experiment. This model predicts the healthiness of muscle cells treated with 30 compounds suspected to induce muscular damage or even myopathies in humans. We also use these profiles to find an interesting morphological similarity between two different classes of drugs: statins (used for cholesterol treatments) and tyrosine kinase inhibitors (anti-cancer drugs). This analysis opens a new perspective for understanding the mechanisms responsible for drug-induced muscular toxicity, an area of toxicology that is currently under-researched.

## Introduction

Despite societal concerns, *in vivo* studies are still a critical step in toxicology as they dictate if a drug candidate is suitable for clinical trials. One of the first building blocks towards the reduction and replacement of animal models is our capacity to evaluate the toxicity potential of drug candidates or potentially dangerous substances using more ethical *in vitro* and *in silico* methods. Predicting the cytotoxicity of a compound such as a small molecule can currently be assessed with well-defined techniques such as QSPR/QSAR (Quantitative Structure-Property/Activity Relationship), which relies on chemical structure descriptors and physicochemical properties of the studied chemical(s) to build models able to predict various endpoints (1). This approach has gained in popularity since the early 2000s, particularly with the introduction of new governmental regulations on all chemicals used in industrial processes, like REACH regulations in the European Union, which requires manufacturers to provide *in silico* toxicology assessment for their products, to identify for instance potential genotoxic, or mutagenic compounds (2).

While this approach has proven that it can accurately predict numerous kinds of drug-induced toxicity such as general cytotoxicity (3), mitotoxicity (4) or even rhabdomyolysis (a fatal myopathy characterized by the breakdown of muscle tissue) (5,6), it is still subject to some limitations. For instance, a QSPR model trained on a chemical family or specific scaffold can still be challenged by an intensively studied phenomenon called the activity cliff, where the observed property (such as toxicity) can sharply decrease/increase with only a slight displacement in the chemical space, and therefore adding complexity to a predictive model (7). Hence, while these QSAR-based models are the backbone of *in silico* toxicity assessment, comparing their findings to complementary approaches would be beneficial for flagging unsuspected dangerous compounds, and for modern drug discovery.

The Cell Painting assay (CPA) is a high-throughput microscopy experiment designed for an extensive evaluation of phenotypic features in fixed cells, using a set of 6 fluorescent biomarkers (8). It allows the simultaneous staining of different organelles of cells in multiple samples to detect morphological changes induced by chemicals. Analysis of the pattern of changes induced by chemicals reveals information about the toxicity of the chemicals and even suggests the chemicals’ mechanism of action (9). The applications of the CPA are diverse, and new ones are continually developed (10,11). Importantly, this approach is independent of chemical structure descriptors and physicochemical properties of the studied chemicals.

While skeletal muscle makes up a substantial portion of human body mass, its toxicity (referred to as myotoxicity) is currently under-investigated: a quick search on Web of Science for research articles dedicated to an organ-specific field of toxicity yields less than 1% of all results for “myotoxicity” (1,141 results), while “neurotoxicity” makes up 43% (65,504 results) (Supplementary Figure 1). In drug discovery and design, the skeletal muscle is usually considered as one of the least important organs for toxicity alerts, which can be due to the relatively low mortality associated with myopathies. This translates into an underrepresentation in clinical trials and research in general (12) with resources allocated to animal studies for other target organs. However, due to its rich blood supply, and the fact that muscle represents 70 to 80% of the human body mass, muscles are highly exposed to circulating drugs (13). Drug-induced myopathies can be serious. For instance, in the late 1990s, the withdrawal of cerivastatin, a lipid-lowering drug used as a treatment for high cholesterol, brought an important problem of rhabdomyolysis (a breakdown of skeletal muscle) to light, detected with post-marketing surveillance (14), with an incidence up to 80 times what was seen in other statins (15). A considerable gap in predictive toxicology research is still to be filled, particularly for rarer toxicities where rapid, inexpensive approaches using *in vitro* models and *in silico* analysis can provide key data. This study aims to address this issue by testing the utility of CPA using a skeletal muscle cell line.

The C2C12 cell line is a mouse myoblast line (16). The cells have been extensively used to study both myogenesis and myotoxicity (17–19). C2C12 cells model a progenitor/stem cell population called satellite cells that are essential for muscle regeneration (20), and can be induced to differentiate into multi-nucleated myotubes (21). The cells therefore provide an *in vitro* model to easily study two different cell types that are central to skeletal muscle biology.

To date, several other studies have assessed cytotoxicity as their prediction endpoint. The predictive potency of phenotypic data extracted from the CPA has already been proven in previous studies, with for instance the prediction of cell heath phenotypes (22) in 3 cell lines, but also various cell health readouts from external assay results taken from public databases such as Tox21 (23).

Here, we used the CPA to create single-cell phenotypic profiles of mouse skeletal muscle cells, following treatment with a panel of thirty reported myotoxicants, at toxicologically relevant concentrations to better understand the responses in cell morphology by building a dataset for myotoxicity prediction in C2C12 cells. The first analysis of our study led us to assess the predictive power of our Cell Painting data gathered from these treatments, to verify if our data contains enough information for this task. Assessment of the predictive power of this newly generated data was performed by creating Random Forest (RF) models towards the prediction of two toxicity endpoints, respectively the cell counts, and the observed cell viability captured with a CTG assay. While recent research projects are witnessing promising results using deep learning methods such as autoencoders (24), RF was here chosen for its ease of use, good model interpretability, high prediction performance, and straightforward feature importance (25). This paper reports on the challenges encountered with the CPA, data collection, data analysis and the utility of this approach for predicting an under researched type of toxicity and makes available a dataset to facilitate future development of the approach.

## Results

### Cell Painting imaging and phenotyping

The CPA revealed a range of morphological and other perturbations induced by toxicant treatment (e.g., Figure 1; Colchicine treatments (COLC)). Phenotypic changes included increased nucleus and cell size (Fig 2A and 2C), decreased count, cells gathered in groups, cell fusion, and important rearrangements in the actin cytoskeleton structure (Fig 2A and 2D). While C2C12 cell fusion to form myotubes is a feature of differentiation, the phenotype of the fused cells was different (Figure 1).

**Figure 1:**
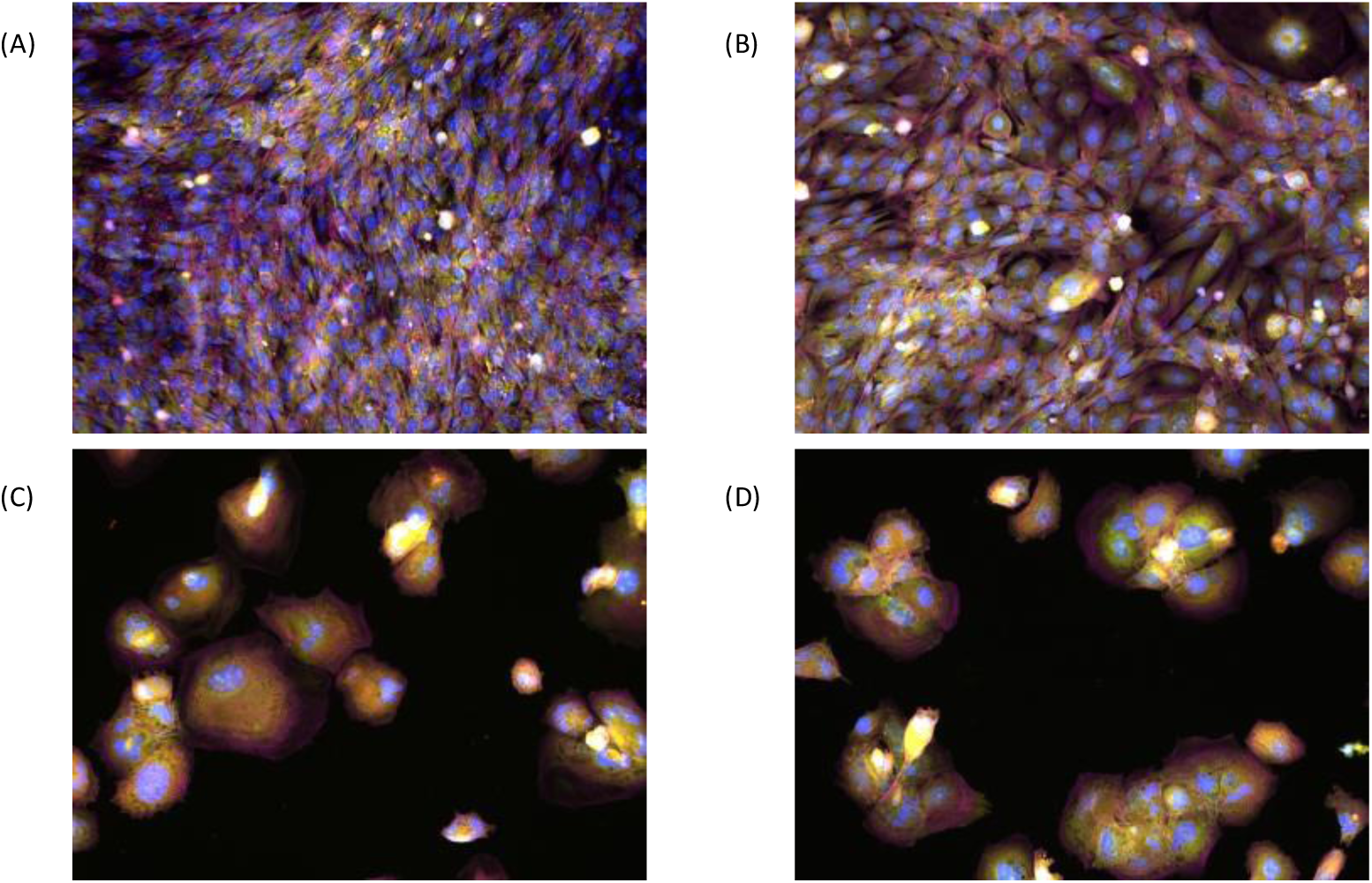
CPA fluorescence microscopy composite images of C2C12 myoblasts captured after a 72 h treatment with DMSO 0.1% as vehicle control (A), or COLC at various concentrations: 10 nM (B), 3 µM (C), 30 µM (D). 20X magnification. Channel color mapping: blue - DNA, red - Mitochondria, green - ER, magenta - Actin, yellow - RNA. Contrast adjusted on DNA channel for improved readability.

**Figure 2:**
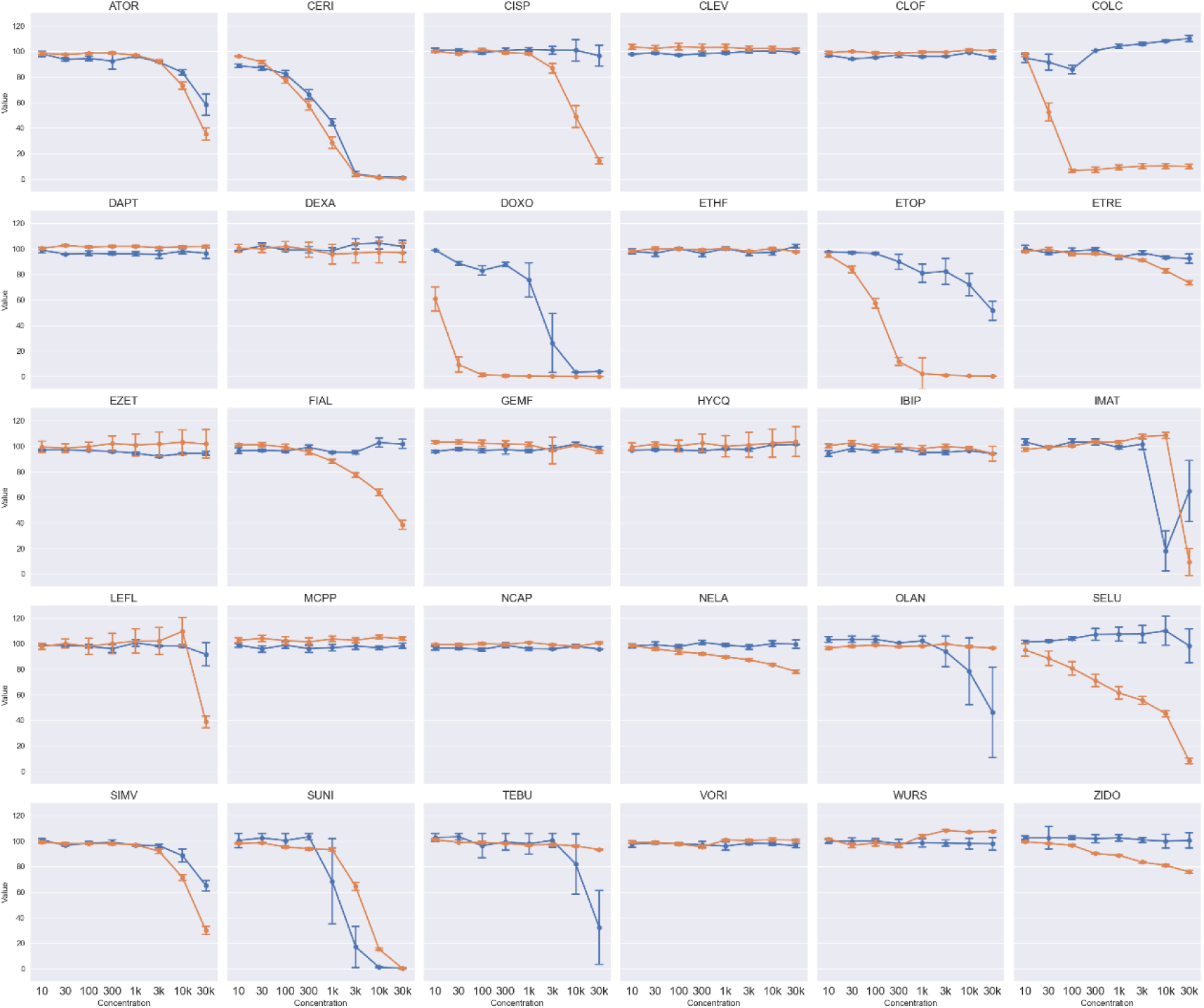
Median cell counts per well (orange lines) and median cell viability (blue lines) plotted against 8 concentrations for the 30 compounds of the Toxifate library in C2C12 myoblasts treatments. Results are shown as median value ± standard error from three biological replicates and four technical replicates, n=12 (3 different plates plated on different days). 5 image fields acquired per well for cell counting.

All the collected images were then processed following an automated protocol, which gave us quantified single-cell phenotypic data for each perturbation, including features describing morphology, intensity, symmetry, and texture properties, totaling 1070 features extracted with Harmony and around 1700 extracted with CellProfiler. A description of feature extraction protocols can be found in Supplementary Table 1. Our CellProfiler pipeline is adapted from the JUMP CPA pipeline for feature calculation, with several tweaks and optimizations, such as typical cell size to match the C2C12 specifications. Comparison of the two feature extraction methods and cell segmentation are beyond the scope of this study and will be discussed in future works. The diversity and number of these toxic phenotypes led us to quantify the changes happening in the cell morphology, starting from the automatization of cell count calculations from these CPA images. The best phenotypes were obtained using the Robust MAD (Median Absolute Deviation) normalization method, as previously recommended by the Carpenter-Singh lab directives and tends to be more popular than a traditional Z-score scaling in recent studies(26–30). These resulting phenotypes were used for further analysis.

### Cell counting from Cell Painting image segmentation

Cell counts were inferred from the analysis protocols, and the resulting median cell counts for all perturbations are summarized in Figure 2 (orange lines). On the common range of concentrations used for this assay, we observed 17 compounds causing a significant decrease in median cell count out of the panel of 30 compared to DMSO controls. Drugs such as ATOR, CERI, SIMV, SELU denote a strong cytotoxic behavior especially in the 3 highest concentrations. Wells treated with SUNI, ETOP or CERI are reaching close to zero detected cells at 30 µM, suggesting that no cell is surviving at this dose, or that their shape and size are not comparable to regular healthy cells. Varying cell sizes can be a challenge for evaluating cell health, which is the reason to compare these findings to an actual cell viability assay.

### Cell viability from an ATP-based assay

Using intracellular ATP measurements from the Cell Titer Glo assay as a surrogate for cell viability is a common approach used to assess toxicant cytotoxicity (31) and is used to derive important points of departure such as NOAEL and LOAEL as well as IC_50_ values that can be valuable in determining the relative toxicity of several compounds.

The results of the three biological replicates of this assay are compiled in Figure 2 (blue lines) and reveal that the most cytotoxic drugs in our panel (in descending order) are CERI, SUNI, DOXO, ATOR, ETOP and IMAT. Several compounds displayed similar trends between cell number and cell count. While the global trend in viability and cell count decrease seems similar between the two assays, it can be noted that the cell count is more sensitive at detecting cytotoxicity, for instance in treatments involving selumetinib (SELU), colchicine (COLC) or etoposide (ETOP). This can be explained by the fact that the CTG assay is only measuring the total ATP content of the wells, which is only monitoring one facet of the actual viability of the cell population.

However, significant discrepancies between cell viability and cell count were observed in many perturbations, with for instance the case of colchicine (COLC), a tubulin-binding gout medication (32). This tubulin disruptor gives the largest difference between these endpoints: while cell count rapidly decreased at 30 nM and then reached a plateau, at the same time only a slight dip in ATP content can be observed, which then started increasing again with higher concentrations according to Figure 2. This striking difference can be attributed to the phenomena of cell fusion triggered by concentrations of COLC at 30 nM and higher. The tubulin disruption caused by COLC forces the actin cytoskeleton to rearrange, and as a stress response, causes the formation of cytoplasmic sacs containing several nuclei, as previously reported in a study employing chick embryo breast muscle (33). It must be noted here that the fusion observed is not related to the usual differentiation of C2C12 myoblasts into myotubes. COLC-induced fusion results in rounder, bigger, and heavily polynucleated cells that do not seem to replicate a myotube-like behavior. The case of FIAL was one of the clearest representations of a decrease in cell count with no effect on ATP content. The Cell Titer Glo signal is constant over the full range of concentrations, while the cell count is steadily declining to reach 40% of the DMSO control value at 30 µM. These findings suggest that relying solely on ATP content might not fully capture the extent of cytotoxic effects, underscoring the importance of considering multiple assays to accurately assess cell health.

### Cell painting phenotypic features can predict cell count

#### Individual models

We next assessed how well the phenotypic profiles extracted with the CPA reflected changes in cell count and cell viability. Starting with the cell count data, a performance summary of 30 RF regression models built with each compound individually is given in Figure 3. The resulting models predicted the median cell count with determination coefficients R^2^ ranging from 0.269 (MCPP), up to 0.996 (CERI), with higher values meaning more accurate predictions. We observed a clear trend in the repartition of determination coefficients, as better performing models were built on compounds showing the most cytotoxic phenotypes, and hence the biggest drop in cell counts with comparison to non-toxic treatments (or controls). In a similar manner, the 5 worst performing models (DAPT, NCAP, EZET, CLEV, MCPP) were built on compounds showing no significative difference in cell number in all tested concentrations. However, such high determination coefficients for all compounds could hide model overfitting, as many values above 0.5 would suggest an important statistical significance of predictions.

**Figure 3:**
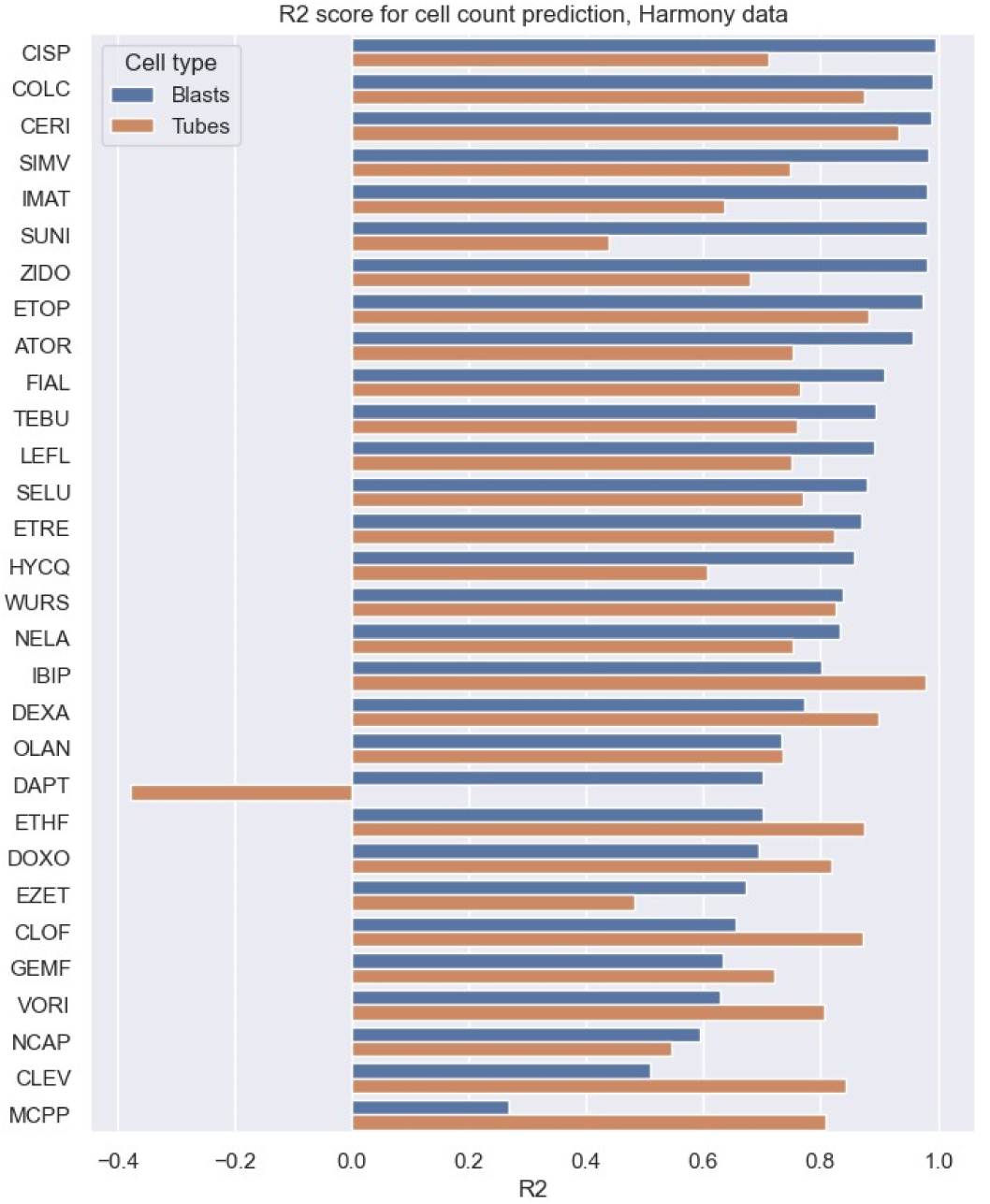
Prediction performance of cell count prediction models built on C2C12 myoblasts and myotubes on Harmony data. Test set determination coefficients (R2) of predicted cell counts against observed cell counts, from Random Forest models built for 30 compounds of the Toxifate library of suspected myotoxicants.

### Cell painting phenotypic features can predict ATP content / cell viability

#### Individual models

Next, we built models predicting cell viability deduced from the Cell Titer-Glo assay. A set of 30 RF models were built on individual compounds using the same approach as described in the cell count prediction methodology. The performance of models followed a trend comparable to the observations in the prediction of cell count (Figure 4), with best-performing models found in the drugs with the most cytotoxic potential, such as CERI, SUNI, and COLC.

**Figure 4:**
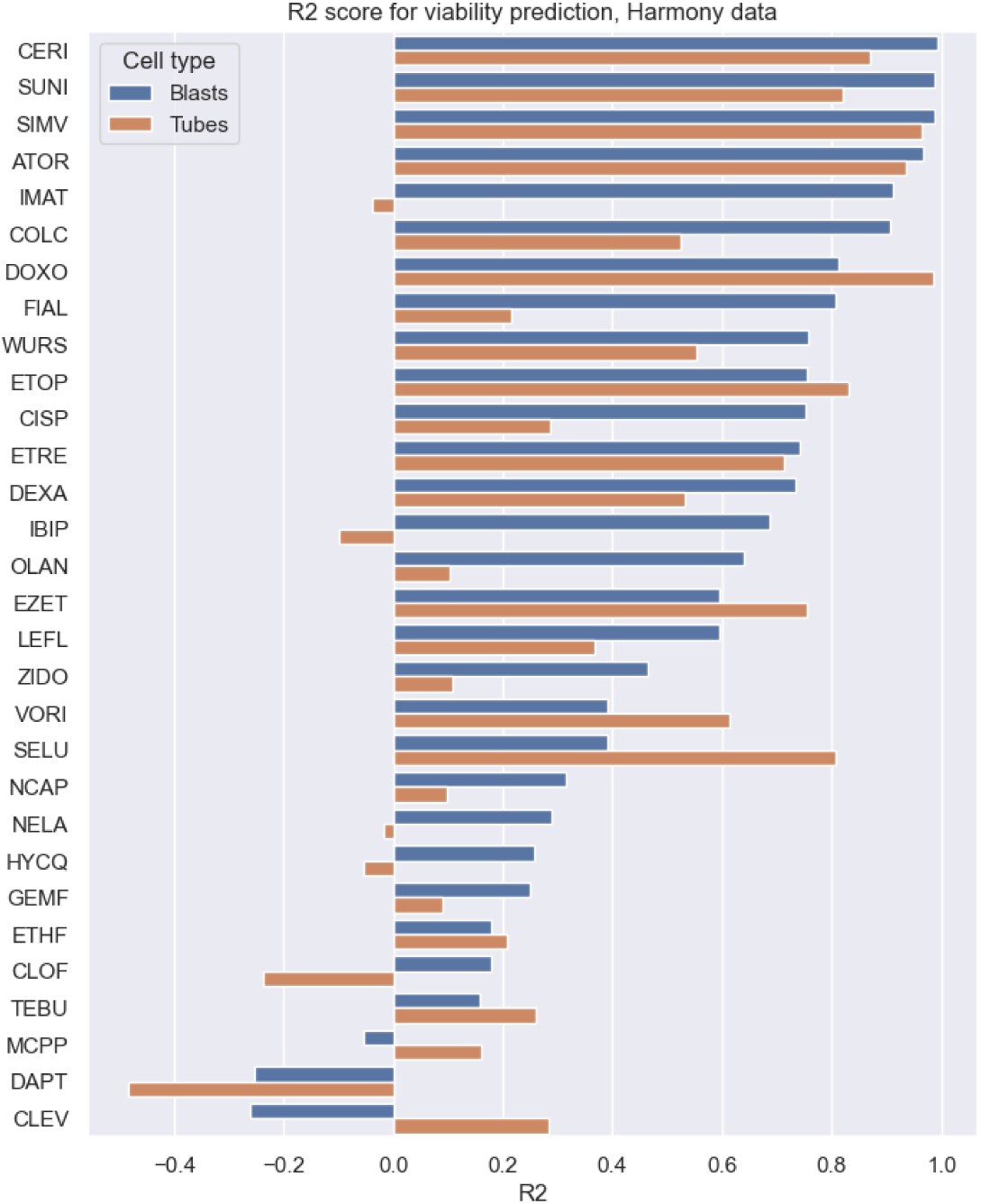
Prediction performance of cell viability prediction models built on C2C12 myoblasts and myotubes on Harmony data. Test set determination coefficients (R2) of predicted cell viability against observed viability, from Random Forest models built for 30 compounds of the Toxifate library. Luminescence data scaled against median luminescence of healthy DMSO controls (represented by 100% luminescence)

A comparison analysis of feature importance for each model showed that individual models of similar compounds yielded similar RF importances, with for instance SUNI and IMAT clustered closely, or also ATOR and SIMV (Figure 5). No clear correlation between the cell viability or counts and the resulting clusters could be seen in this analysis.

**Figure 5:**
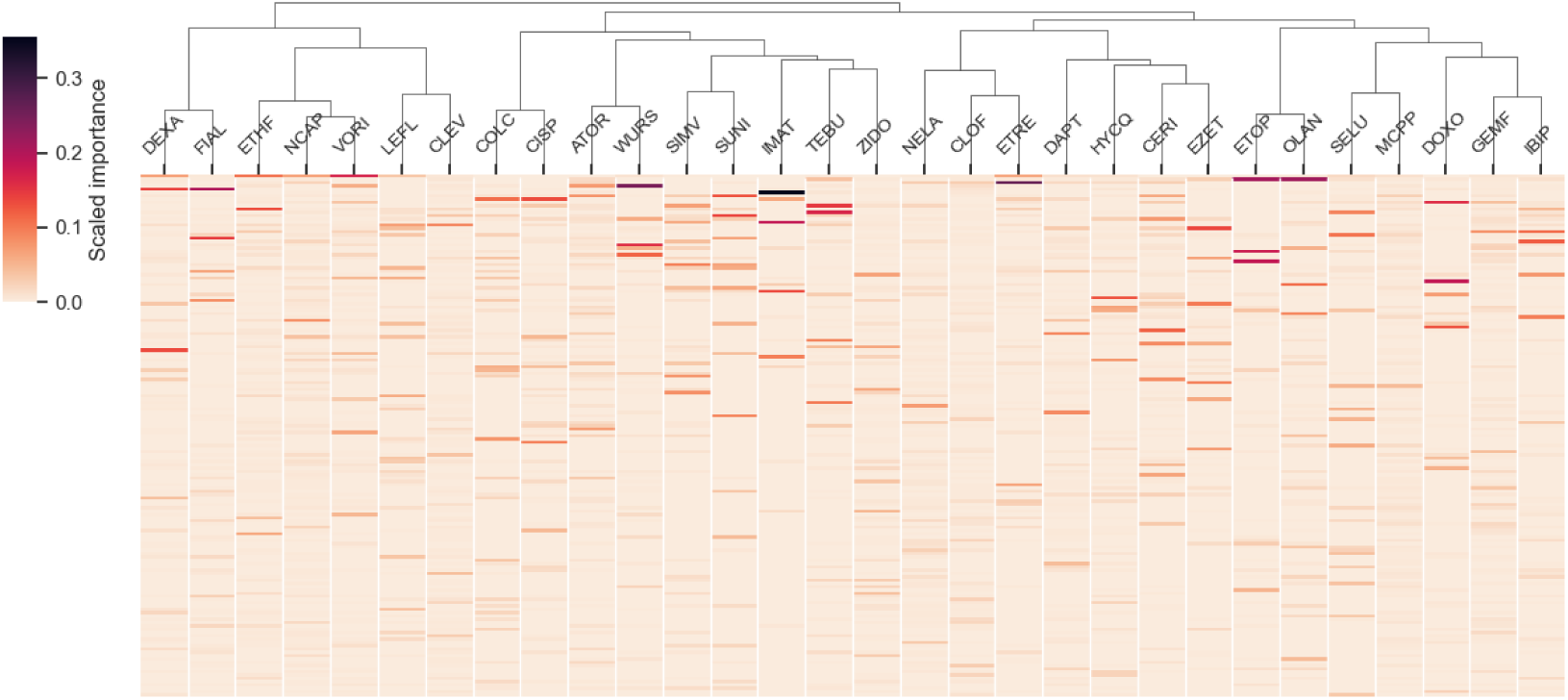
Heatmap and hierarchical clustering of Random Forest feature importance for each model built on individual drugs for viability prediction in myoblasts. Features are ranked in descending order by the sum of their scaled importance across all models (most important features are at the top of the heatmap). Clustering is performed using single-linkage and correlation metric. The lower half of the heatmap containing less prominent features was cropped for readability.

#### RF model trained with all compounds

A single RF model was then trained on the entire drug panel on myoblast data, using well-level profiles. The dataset is split into a training and a test set (80-20 %), and yields a test set R^2^ of 0.861, and a mean squared error of 59.8, indicating that this model can predict cell viability with good accuracy (training set: R^2^ = 0.983, MSE = 6.67). The 10 most important features (according to mean decrease in impurity) of the RF model are shown in Table 2. An assessment of feature importance in the resulting RF shows that cytoplasm-related features are prevalent for cytotoxicity prediction, especially for Spots Edges and Ridges (SER) Profiles in the ER and mitochondria channels “ ytoplasm Mito Profile 4/5 SER-right” and “ ytoplasm 488 Profile 4/5 SER-Edge”. The feature describing the Nucleus profile in the Hoechst N channel “Nucleus 33342 Profile 5/5” was also found to be crucial in several models.

**Table 2:**
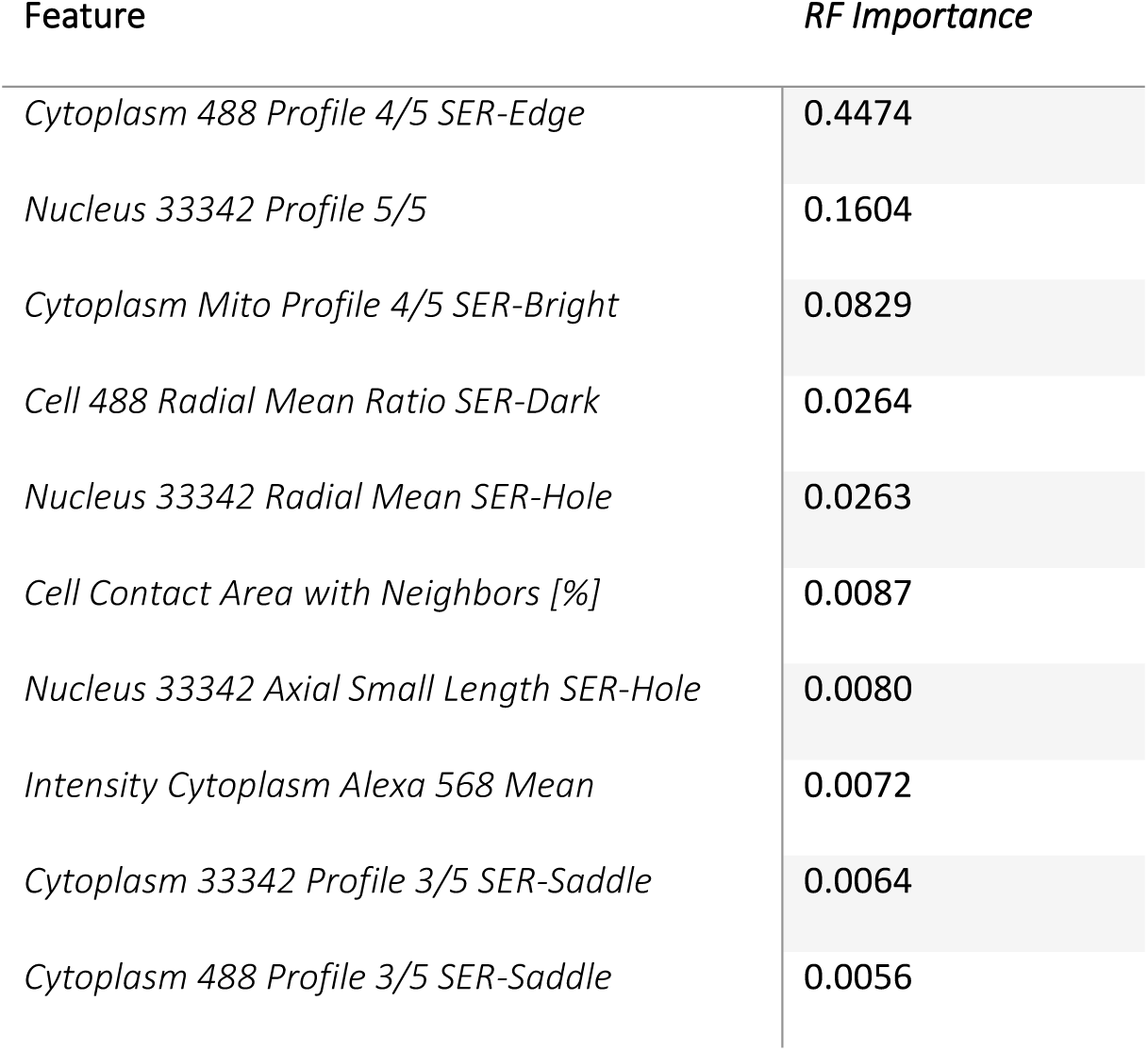
Summary of RF feature importance for the top 10 properties of the Random Forest model for viability prediction built on C2C12 myoblasts.

#### Prediction of viability and cell count in myotubes

The same approach was followed for predicting cell viability and cell count in C2C12 differentiated myotubes, using the same segmentation algorithm in Harmony 4.8 as for myoblasts (Figure 3 and Figure 4). Model performance followed the same trends as the myoblasts, with better predictions given for most cytotoxic compounds, and less reliable models built on compounds displaying no significant signs of toxicity, hence models are capturing noise in the cell viability and cell count. No discrepancy in descriptor importance was noted, as the same feature classes were found with comparable importance.

### Cell Painting phenotypes can uncover unsuspected similarities between the drugs

To obtain a broader representation of the biological information encoded in each quantitative phenotype, and of the relationships between different cytotoxic perturbation, a clustering of the normalized profiles was conducted, and the resulting clusters and profiles heatmap of the myoblasts is shown in Figure 6. Morphological changes are considered minor when they display an absolute Z-score values below three, which allows us to keep only the relevant phenotypes containing major and significant perturbations. In Figure 6 are represented myoblasts profiles displaying an induction greater than 0.2, meaning that at least 20% of features in that phenotype are significantly affected. This induction threshold (20% affected features or greater) used for filtering out low-cytotoxic perturbations was found to be efficient on our C2C12 data and excluded profiles with minor morphological changes in myoblasts and myotubes, that could have spoiled the clustering with meaningless clusters.

**Figure 6:**
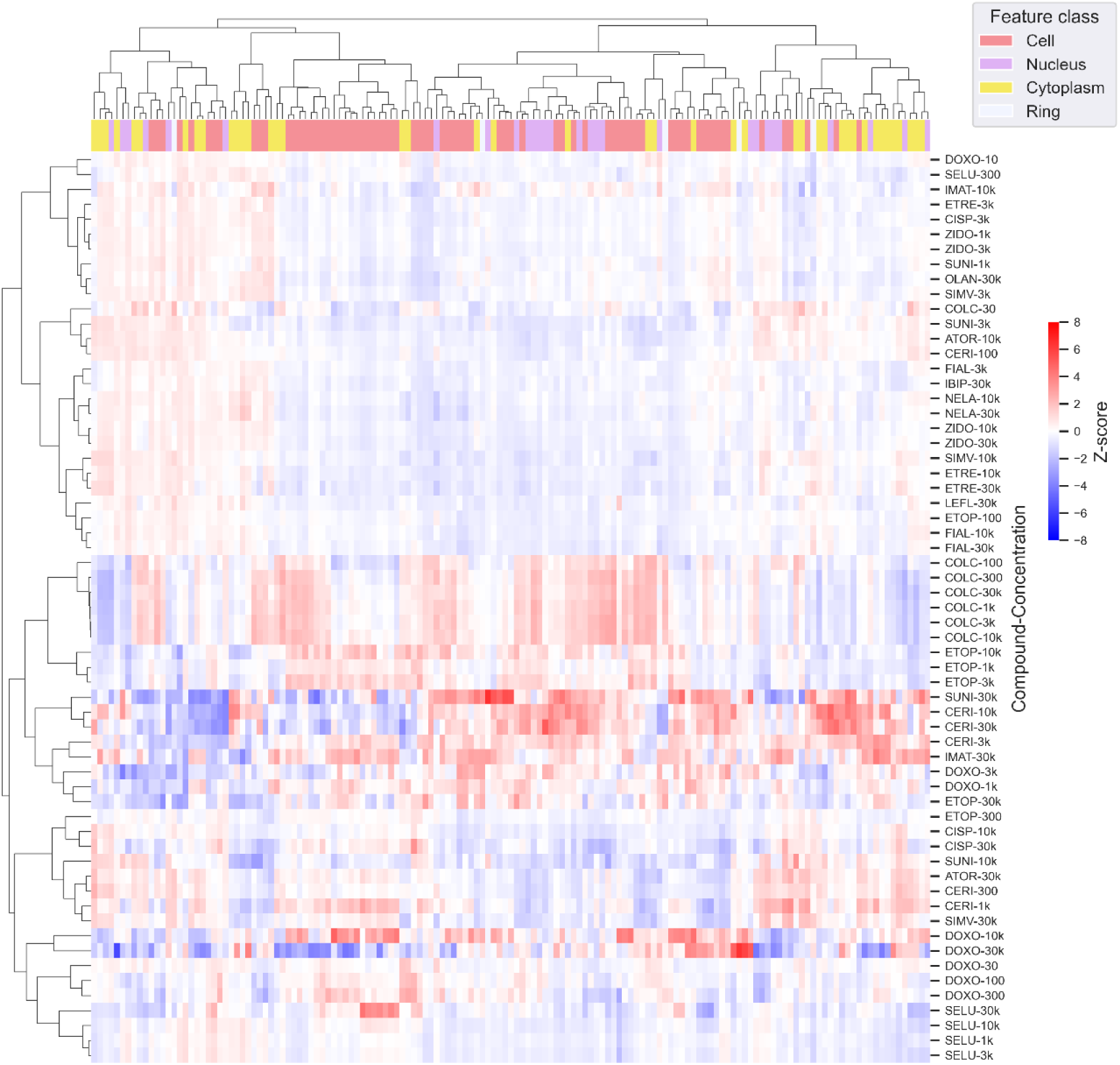
Heatmap and hierarchical clustering of treatment-level morphological profiles for C2C12 myoblasts. Only profiles with induction > 0.2 are represented. Hierarchical clustering of treatments and features computed with complete-linkage on Pearson correlations. Z-scores clipped to ±8 MAD for increased readability and color scaling.

The distribution of induction values for the two cell types is given in Supplementary Figure 2. It was found that the hierarchical clustering using average linkage method and a correlation metric between profiles yielded the most satisfactory results with distinct clusters of toxic phenotypes. Because of the high dimensionality of the data only a few distance metrics are suitable for that type of vector. Euclidean distance is here yielding poor results due to the “curse of dimensionality” (34). A Mahalanobis distance would have been efficient by mixing numerical distance and correlation, but it was limited by the total number of features being greater than the number of samples (as usual in cell phenotype analysis), making its use impossible.

Several expected clusters are emerging from this analysis. For instance, statins tend to group together, even at different concentrations. ATOR-30k, CERI-300, CERI-1k, SIMV-30k, are all displaying remarkably similar profiles, and this observation is not surprising as these 3 drugs are members of the statin family, being 3-hydroxy-3-methylglutaryl-CoA reductase (HMGCR) inhibitor. These samples are hence clustered in accordance with their biological target, while their chemical structure is however more dissimilar, and do not follow a clear scaffold. More surprisingly, in the three statin clusters (one with low cytotoxicity, one moderate and one displaying advanced cytotoxicity), we observe the presence of SUNI, a tyrosine kinase inhibitor (TKi) which has no structural similarity to statins and has a different pharmacological target. Previous studies have shown the cytotoxic potential of SUNI in C2C12 cells (35). Several cases of SUNI induced rhabdomyolysis have been reported in the past (36), which is one of the most serious adverse effects observed globally with statins, and many lipid-lowering drugs. Thus, it appears that statins and SUNI display a similar cytotoxicity in C2C12 cells and in muscle, although the underlying mechanisms that explain this similarity are unclear. To further investigate the relationship between sunitinib and the statin family, a connectivity search on gene expression data was conducted on cMAP (37), accessed from the clue.io platform. A comparison of sunitinib with the 3 statins from our panel displayed many similarities on the perturbagen class and on targets. Sunitinib appears to be a HMGCR inhibitor, or at least share similarities in gene expression, but also shares the properties of a proto-oncogene tyrosine-protein kinase (SRC) inhibitor such as dasatinib, and of a norepinephrine reuptake inhibitor, also displayed by the statins. The genetic connectivity was less conclusive, with only a few genes of interest targeted by the four compounds, like PCSK9 (Proprotein convertase subtilisin/kexin type 9) a target for the treatment of dyslipidemia, or MED4 (mediator of RNA polymerase II transcription subunit 4, also known as DRIP36). Another cluster of interest is standing out in this analysis: the proximity of NELA and zidovudine ZIDO at concentrations of 10 and 30 µM is observed in C2C12 myotubes and in myoblasts, suggesting a similarity in toxic mode of action. NELA is commonly used as a treatment for T-cell acute lymphoblastic leukemia, while ZIDO is used as an HIV treatment and preventive medication, targeting the HIV-reverse transcriptase (38). Both compounds are nucleoside analogues and share the inhibition of DNA synthesis as a common mechanism of action. Structure-wise, ZIDO is a thymidine analogue, with an azide substituent replacing the cyclic alcohol group. NELA is a purine nucleoside analogue with an additional OH group added to the sugar ring, and a methoxy group replacing the cyclic ketone.

## Discussion

The CPA conveys meaningful information on many aspects of cell health and cell viability in cell lines (22). This study confirms its ability to predict these endpoints, even when working with more specific cell line models like C2C12. The main limitations of this study include the small size of the drug panel, containing only 30 compounds, which is low for usual predictive toxicology standards. However, the small panel size is counterbalanced by testing each chemical at a range of concentrations, allowing detection of concentration dependent differences in phenotypes. For instance, Imatinib is following a trend of morphological perturbations similar to simvastatin on most concentrations but appears vastly different at 300 nM, and then even gets close to a high dose cerivastatin-like phenotype at 30 µM. These findings highlight the importance of choosing the “right” concentration for testing a single dose in other CPA. A common approach would be to use the LOAEL or NOAEL (lowest or no observed adverse effect level) for each compound. To overcome the effect of concentration on changing phenotypes and compare these with single-dose gene expression data for instance, one solution is to use a technique such as concentration-response modelling to find a “point of departure” for each feature category, as detailed by Nyffeler et al. (26,39). However, keeping a single value for each compound could omit valuable information on potentially unsuspected pathways activated at specific concentrations.

It can be pointed out that the dataset used in this study is quite unbalanced, as many treatments have little to no statistically significant effect on cell viability or cell count, leading to an underrepresentation of cytotoxic morphological profiles. To try overcoming this problem, a synthetic data generation algorithm such as SMOGN was used, to compensate the unbalanced distribution of viabilities(40). This algorithm is the generalization to regression tasks of the well-known SMOTE generative method, dedicated for classification models (41). Promising results were obtained using the Python implementation, with the generation of new cytotoxic profiles with gaussian noise applied to morphological features, and results are shown in Supplementary Figure 3. The ideal case is however to modify the experimental approach, to account for more cytotoxic profiles, for instance by including compounds with fewer reports linked to toxic myopathies, and also avoiding compounds with no visible effect on cell morphology, or no effect on viability.

The implementation of this high-content microscopy assay raised important challenges on diverse aspects of the project. Crosstalk between the ER and the RNA channels has been observed regularly, reducing the total amount of information conveyed by the 5 channels. This is however not an exception as such an excitation bleed through can be seen in other CPA studies (42). The DNA, actin, and mitochondria channels were not affected by this crosstalk, as their respective excitation wavelengths are far apart enough. Moreover, in this study, a 20 × long air objective was used in wide field mode, as opposed to the more popular 20 × water immersion objective in confocal imaging, which captures more signal from the excited fluorophores, and provides higher quality images compared to the air objective. However, we proved that even with non-optimal conditions phenotypic changes are still captured effectively, and a clear signal is extracted. In several cases, significant discrepancies between the cell viability of a treatment and the corresponding cell count were seen. For instance, a treatment of C2C12 myoblasts with cisplatin seems to have an important impact on the number of cells detected in a treatment well, whereas ATP content of the cell population will not be significantly modified.

It is to be noted that the choice of the right model of microscopy plate is of utmost importance for this kind of assay, especially when cell adherence is a crucial factor, as we noticed that myotube shape was different between plate models. Slight differences in plate specification, manufacturer, or even manufacturing batch can lead to significant phenotypic differences. This phenomenon was mainly observed when capturing myotube data (as their shape is very dependent on the well coating). Mitigation is possible with rigorous normalization, including techniques such as Robust MAD, and B-score plate normalization (43).

Regarding the phenotypic clustering of perturbation profiles, the idea of implementing connectivity constraints to the hierarchical clustering surely would reduce the occurrence of some compounds in multiple clusters, seen with a compound such as ATOR not creating a single cluster. However, these constraints may obscure the diversity of the toxicity profiles for this kind of drugs.

It will be compelling in following studies to pinpoint specific features or features groups in the morphological phenotypes to correlate precise changes in morphology and stress signaling pathways or early signs of cytotoxicity, as already introduced by a few studies to date (23). This next step will allow us to capture all the necessary information to potentially describe and predict differential gene expression, or more broadly pathway activation and inhibition. The CPA is designed in a context of high-content microscopy, which usually involves the allocation of substantial resources, especially with a team of diverse scientists in each step of the assay pipeline, from the cell culture to the data analysis. However, another examples have shown the accessibility of this experiment in smaller chemical biology laboratories (44). In future works, the differences in the feature calculation and segmentation methods used in this study will be assessed. Moreover, a comparison of drug profiles obtained in the C2C12 cell line will be compared with data published in other cell lines, especially with U2OS used by many image-based profiling teams.

To conclude, this high-content microscopy assay is categorically promising for bridging gaps in toxicological data and brings another point of view to ensure reducing the need for *in vivo* assays. This study is, to our knowledge, the first occurrence in literature of an application of the CPA to the C2C12 cell line, moreover to its differentiated state, proving the versatility and adaptability of this assay to new cellular systems.

## Materials and methods

### Drug library

A collection of 30 myotoxicants (Table 1) was prepared in a 96-well plate format. The toxicants were dissolved in DMSO (or DMF for cisplatin) at a range of 8 concentrations from 10 µM to 30 mM and three replica plates were prepared. The plates were sealed and stored at −20 °C until use. Plates were thawed at room temperature in the dark for at least 2 hours before use.

**Table 1:** Toxifate drug panel of myotoxicants.

| **Drug name** | **Class** | **Abbreviation** | **Supplier** | **ChEMBL ID** |
| --- | --- | --- | --- | --- |
| *Aminocaproic acid* | Anti-fibrinolytic | **NCAP** | Sigma | CHEMBL1046 |
| *Atorvastatin* | Lipid lowering | **ATOR** | Sigma | CHEMBL1487 |
| *Cerivastatin* | Lipid lowering | **CERI** | Sigma | CHEMBL1477 |
| *Cisplatin* | Chemotherapy | **CISP** | Sigma | CHEMBL11359 |
| *Clevudine* | Hepatitis B | **CLEV** | Santa Cruz Biotech | CHEMBL458875 |
| *Clofibric acid* | Lipid lowering | **CLOF** | Sigma | CHEMBL683 |
| *Colchicine* | Gout | **COLC** | Sigma | CHEMBL107 |
| *Daptomycin* | Antibiotic | **DAPT** | Sigma | CHEMBL387675 |
| *Dexamethasone* | Corticosteroid | **DEXA** | Sigma | CHEMBL384467 |
| *Doxorubicin* | Chemotherapy | **DOXO** | Cayman Chemicals | CHEMBL53463 |
| *Ethyl fluoroacetate* | Rodenticide | **ETHF** | Sigma | CHEMBL509273 |
| *Etoposide* | Chemotherapy | **ETOP** | Sigma | CHEMBL44657 |
| *Etretinate* | Retinoid | **ETRE** | Sigma | CHEMBL464 |
| *Ezetimibe* | Lipid lowering | **EZET** | Sigma | CHEMBL1138 |
| *Fialuridine* | Hepatitis B | **FIAL** | Sigma | CHEMBL271475 |
| *Gemfibrozil* | Lipid lowering | **GEMF** | Sigma | CHEMBL457 |
| *Hydroxychloroquine* | Anti-malarial | **HYCQ** | Sigma | CHEMBL1535 |
| *Ibipinabant* | Cannabinoid | **IBIP** | Santa Cruz Biotech | CHEMBL412262 |
| *Imatinib* | Chemotherapy | **IMAT** | Sigma | CHEMBL941 |
| *Leflunomide* | Anti-rheumatic | **LEFL** | Sigma | CHEMBL960 |
| *Mecoprop* | Herbicide | **MCPP** | Sigma | CHEMBL272942 |
| *Nelarabine* | Chemotherapy | **NELA** | Sigma | CHEMBL1201112 |
| *Olanzapine* | Antipsychotic | **OLAN** | Sigma | CHEMBL715 |
| *Selumetinib* | Chemotherapy | **SELU** | Santa Cruz Biotech | CHEMBL1614701 |
| *Simvastatin* | Lipid lowering | **SIMV** | Cayman Chemicals | CHEMBL1064 |
| *Sunitinib* | Chemotherapy | **SUNI** | Cayman Chemicals | CHEMBL535 |
| *Tebuconazole* | Antifungal | **TEBU** | Sigma | CHEMBL487186 |
| *Voriconazole* | Antifungal | **VORI** | Sigma | CHEMBL638 |
| *Wurster's blue* | Redox indicator | **WURS** | Sigma | CHEMBL1393325 |
| *Zidovudine* | Antiretroviral | **ZIDO** | Sigma | CHEMBL129 |

### C2C12 cell culture

Proliferating C2C12 myoblasts (Sigma-Aldrich, CRL-1772) were cultured in a solution of Dulbecco’s Modified Eagle Medium (DMEM) (Sigma) supplemented with 20% Fetal Bovine Serum (FBS) (Sigma) and an antibiotic cocktail of 1% Penicillin and Streptomycin (PS) (Sigma Aldrich). Cells were passaged at 60-70% confluence. Cells to be differentiated into myotubes were seeded at 5,000 cells per well into 384 wells plates (Greiner Bio-One µClear) using a BioTek Multiflo automated dispenser at 20 µL per well. Cells were cultured for 4 days in a differentiation medium (DM; DMEM supplemented with 2% Horse Serum (Sigma) and 1% PS) before being treated with drugs. Myoblasts were seeded at 1,000 cells per well on day 4 in a growth medium (GM; DMEM supplemented with 20% Fetal Bovine Serum and 1% PS).

### Drug treatment

The two C2C12 cell types were treated on day 4 using a PerkinElmer JANUS automation system for optimal experimental reproducibility. Cells were treated with toxicants at 8 different concentrations from 10 nM to 30 µM. The final concentration of vehicle (DMSO for all drugs except cisplatin, which was dissolved in DMF) was 0.1%. Plates were then incubated at 37°C and 5% CO_2_ for 72 hours in the dark. Three biological replicates run on separate days were completed with 4 technical replicates of each condition on each plate, over 5 well sites. Each perturbation (one drug at one concentration) was hence repeated in 12 wells to capture a large population of cell phenotypes.

### Cell Painting staining and image acquisition

At 72 hours, the cells were fixed and stained following the JUMP CPA protocol from Carpenter lab (Figure 7), using the JANUS automated dispensing platform. Plates were then imaged using a Perkin Elmer Operetta equipped with a 20 × long air objective. Five fields were captured for each well, with no pixel binning. Images were analyzed with Harmony 4.8 following the steps described in Nyffeler et al. (26), and then also analyzed externally using the open-source Cell Profiler software (45). A summary of this protocol is outlined in Figure 1.

**Figure 7:**
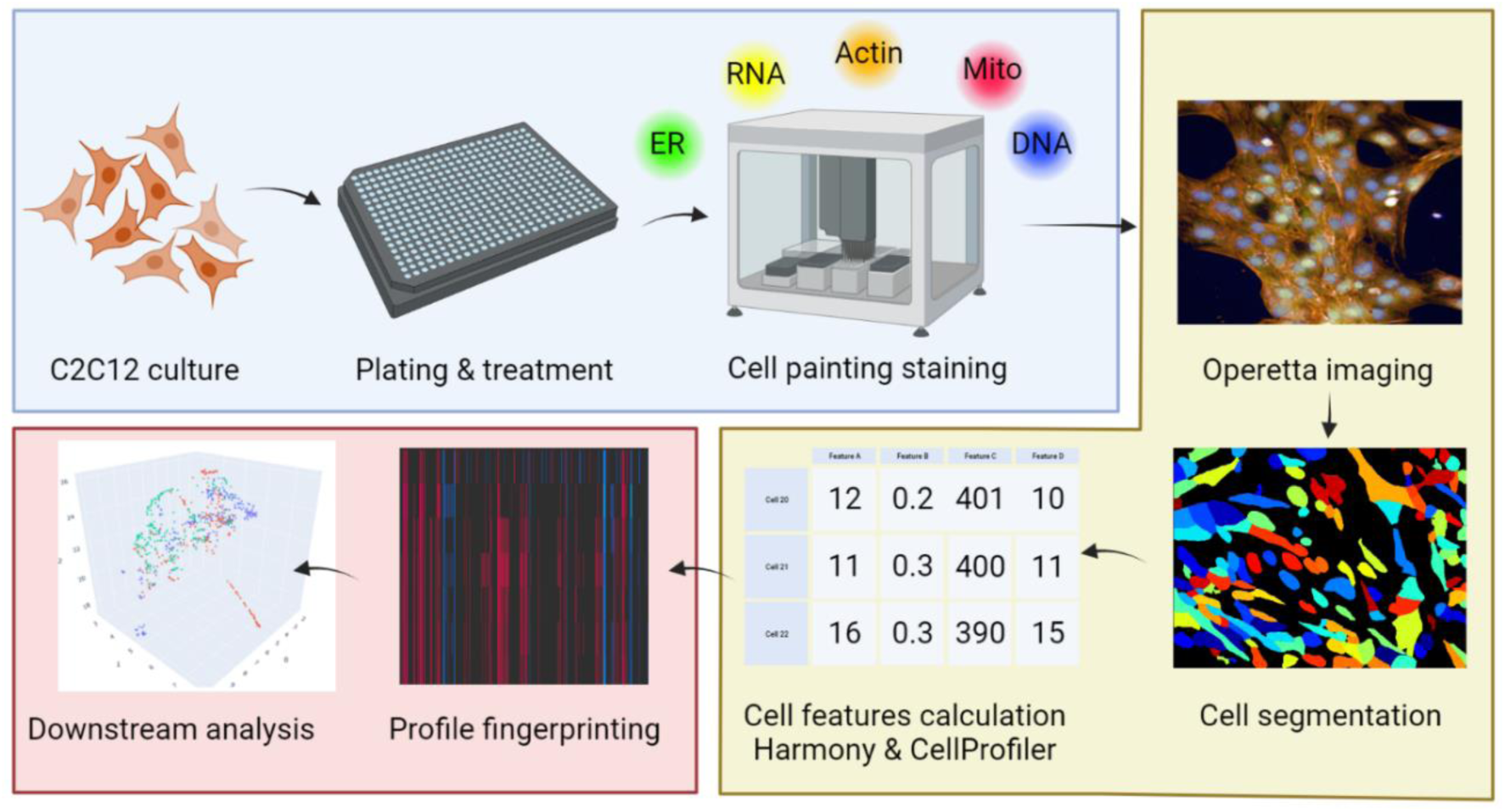
Outline of the CPA protocol applied to C2C12 cells. Color coding, Blue: cell culture and staining, Yellow: high-content microscopy acquisition, Red: data analysis

### Data analysis

Data analysis was performed in KNIME 4.7 and Python 3.10 with the pycytominer package for per-well profile aggregation and normalization (46). Normalization methods used were min-max scaling, Z-score, and Robust MAD, as implemented in pycytominer. The well-level phenotypes were then aggregated from the single-cell data, by taking the median feature values for each well population, and then normalizing on the negative vehicle control wells (30 to 40 wells per plate) with a Z-score method. Cell counts per well were also calculated for comparison with the cell viability assay.

### Profile clustering

Perturbation-aggregated profiles were clustered using hierarchical clustering, with complete linkage method. The value of induction of a well-level profile was computed as the percentage of morphological features with an absolute Z-score value greater than 3, as introduced by Schneidewind et al (9) :

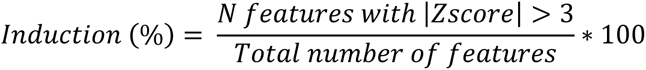

This metric allows a simple filtering of perturbations with low phenotypical significance and high noise.

### Cell viability assay

The same procedure for cell culture and drug treatment as the CPA was followed for assessing cellular viability with ATP content. After 72 h of treatment, 20 µL of Cell Titer-Glo reagent (Promega, Walldorf, Germany) was added to samples, plates were incubated in the dark at room temperature for 30 minutes, and luminescence was read with a Perkin Elmer VICTOR3 plate reader. Data was exported as an XLS file containing raw luminescence reads, which was then normalized, and expressed as a percentage of viability of each plate vehicle controls (DMSO).

### Viability prediction model building

The median cell count per well was calculated with the number of valid objects segmented during image analysis, processed with a custom imaging pipeline set up in Harmony. Phenotypes pre-processing and model building are conducted in Python 3.10 and sklearn (47). Features presenting standard deviation higher than 2,000 were discarded, to remove potentially large values caused by the normalization process. Correlated features were also removed with a threshold of 0.9. A RF regression model (25) was built for each toxicant (excluding DMSO controls) using all remaining features, and each model was validated by a test set using random sampling of well-level phenotypes (train test ratio of 80/20). 96 observations were hence studied for each toxicant (8 concentrations × 4 wells per plate × 3 plate replicates).

## Acknowledgements

Special thanks to Enda O’ onnell for the supervision and help brought through the Screening core facility of the University of Galway for fluorescence microscopy and liquid handling automation with the JANUS platform.

## Data availability statement

Morphological profiling data obtained in C2C12 can be openly accessed on the dedicated Zenodo repository of this article: DOI 10.5281/zenodo.10623852. Due to the extensive size of single cell profiling data and associated TIFF images, this data is available on demand to the corresponding author.

## Code availability

Jupyter Notebooks, Python scripts and KNIME workflows used in this study are accessible from the project GitHub repository at the following address: https://github.com/romlambert/CytotoxPred

## Supporting information

**Supplementary Table 1:** Summary of extracted features with Harmony and CellProfiler software. (A) Features extracted via Harmony software, not including cell positional features and basic morphology, following guidelines from Nyffeler et al (26). (B) Summary of features extracted with CellProfiler. N = nuclei; Ce = cell; Cy = cytoplasm; M = membrane; R = ring region, x=all compartments. Cells are segmented into 5 compartments in Harmony, and in 3 with CellProfiler (nuclei, cytoplasm, and cell)

| Channel | Symmetry | Compactness | Axial | Radial | Profile | Intensity | Texture |
| --- | --- | --- | --- | --- | --- | --- | --- |
| DNA | N | N | N | N Ce | N Cy | N | N |
| RNA | N | N | N | N | N | N | N |
| ER | Ce | Ce | Ce | Ce | Cy | R Cy | R Cy |
| AGP | Ce | Ce | Ce | Ce | N Cy | R Cy M | R Cy M |
| MITO | Ce | Ce | Ce | Ce | N Cy | R Cy | R Cy |

**(B)**
| Channel | Colocalization | Granularity | Intensity | Neighbors | Intensity distribution | Size & shape | Texture |
| --- | --- | --- | --- | --- | --- | --- | --- |
| DNA | X | X | X | X | X | X | X |
| RNA | X | X | X | X | X | X | X |
| ER | X | X | X | X | X | X | X |
| AGP | X | X | X | X | X | X | X |
| MITO | X | X | X | X | X | X | X |

**Supplementary Table 2:** Connectivity results of sunitinib against the statins ATOR, CERI, SIMV obtained from the cMAP tool ([clue.io]). cMAP *tau* scores of top perturbagen classes (A), top genes (B), and top compound similarities (C). Scores ranging from −100 to +100, and values over 90 or below −90 are considered of interest for further investigation. Values outside this threshold are outlined in red. Only compound BRD-M64432851 was considered for sunitinib due to the presence of duplicates. Duplicate compound names were truncated for readability.

**(A)**
| Perturbagen class | score | ATOR | CERI | SIMV | SUNI |
| --- | --- | --- | --- | --- | --- |
| HMGCR inhibitor | 99.96 | 100.00 | 99.95 | 99.97 | 91.51 |
| PI3K inhibitor | 99.14 | 99.11 | 59.41 | 99.36 | 99.16 |
| DNA dependent protein kinase inhibitor | 98.92 | 98.93 | 62.09 | 99.35 | 98.91 |
| SRC inhibitor | 98.76 | 99.53 | 95.10 | 99.48 | 98.04 |
| Serotonin receptor antagonist | 98.70 | 99.14 | 90.67 | 99.48 | 98.27 |

**(B)**
| Gene symbol | score | ATOR | CERI | SIMV | SUNI |
| --- | --- | --- | --- | --- | --- |
| MED4 | 95.99 | 95.92 | 96.05 | 96.98 | 81.95 |
| OVOL2 | 94.71 | 98.25 | 93.50 | 95.91 | 86.16 |
| C9ORF96 | 94.39 | 98.70 | 94.69 | 94.09 | 81.47 |
| MST1R | 94.20 | 98.89 | 65.27 | 97.74 | 90.67 |
| GPR110 | 93.86 | 98.66 | 14.33 | 95.53 | 92.20 |
| ATP6V1D | 93.15 | 96.90 | 86.54 | 95.07 | 91.23 |
| PCSK9 | 92.88 | 94.71 | 93.60 | 92.15 | 81.86 |
| RXRA | 92.79 | 94.55 | 8.34 | 91.03 | 94.96 |

**(C)**
| Compound name | score | ATOR | CERI | SIMV | SUNI |
| --- | --- | --- | --- | --- | --- |
| RS-17053 | 99.83 | 99.86 | 92.47 | 99.86 | 99.79 |
| BIBU-1361 | 99.77 | 99.82 | 99.49 | 99.79 | 99.75 |
| wortmannin | 99.65 | 99.68 | 87.96 | 99.82 | 99.61 |
| fluvastatin | 99.58 | 99.89 | 99.30 | 99.86 | 88.73 |
| lovastatin | 99.56 | 99.89 | 99.33 | 99.79 | 85.88 |
| NNC-55-0396 | 99.56 | 99.30 | 71.81 | 99.82 | 99.82 |
| CGP-71683 | 99.53 | 99.44 | 61.99 | 99.79 | 99.61 |

**Supplementary Figure 1:**
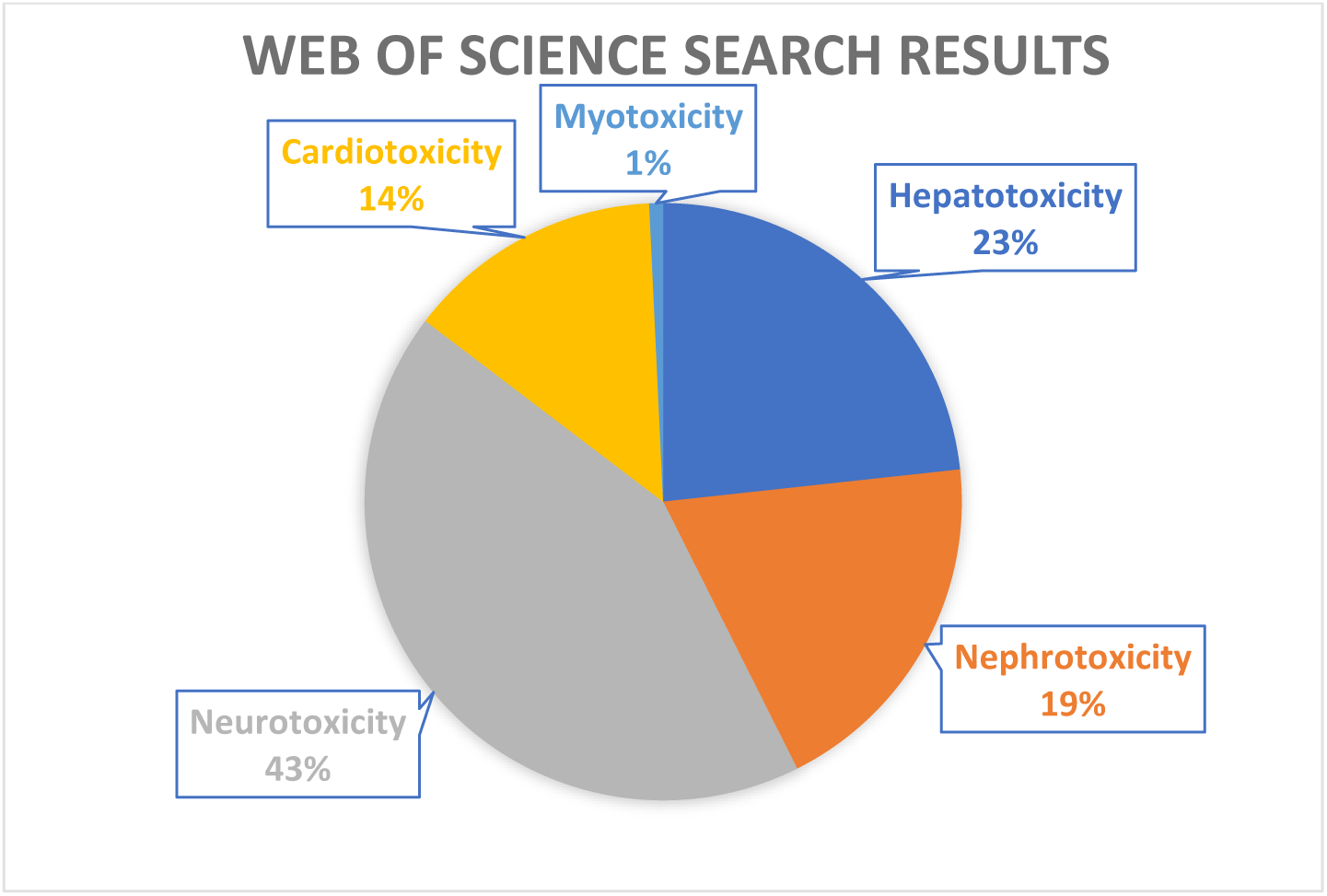
Results of a document search on the Clarivate Web of Science platform performed on August 1^st^ 2023, in percentage of total documents yielded with all queries. Keywords were entered as displayed in lowercase, and no synonyms were considered. Search conducted in the WoS Core Collection only, including all document types.

**Supplementary Figure 2:**
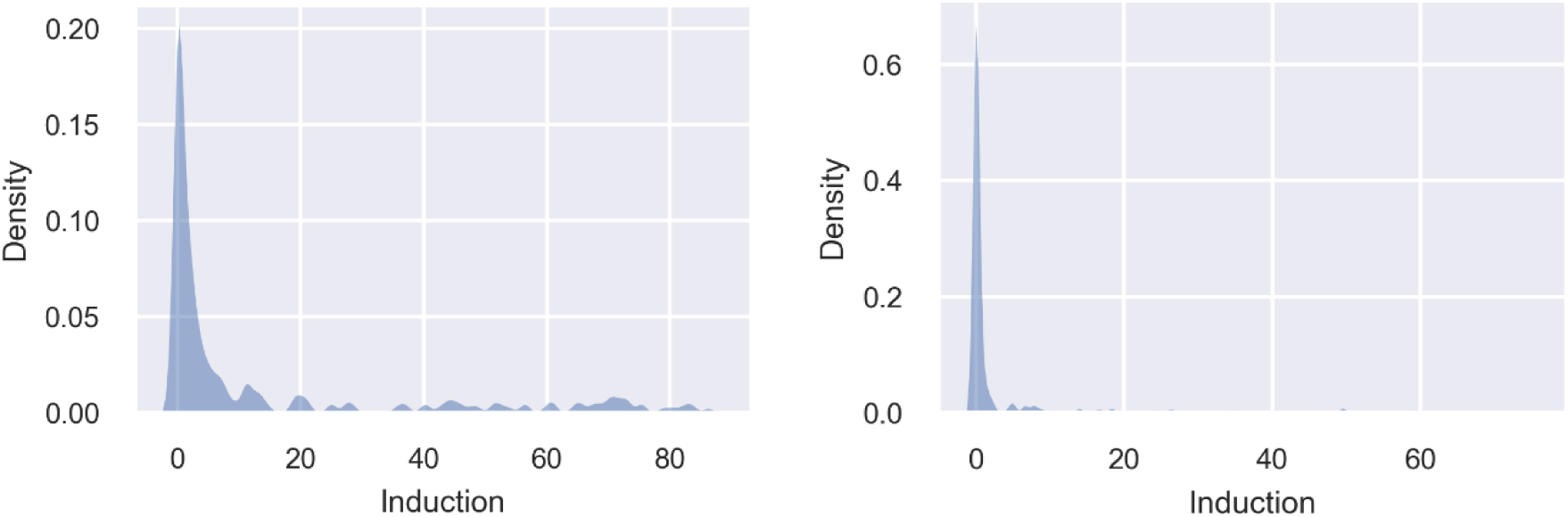
Distribution of induction values for myoblasts (left) and myotubes (right) treatment-level data.

**Supplementary Figure 3:**
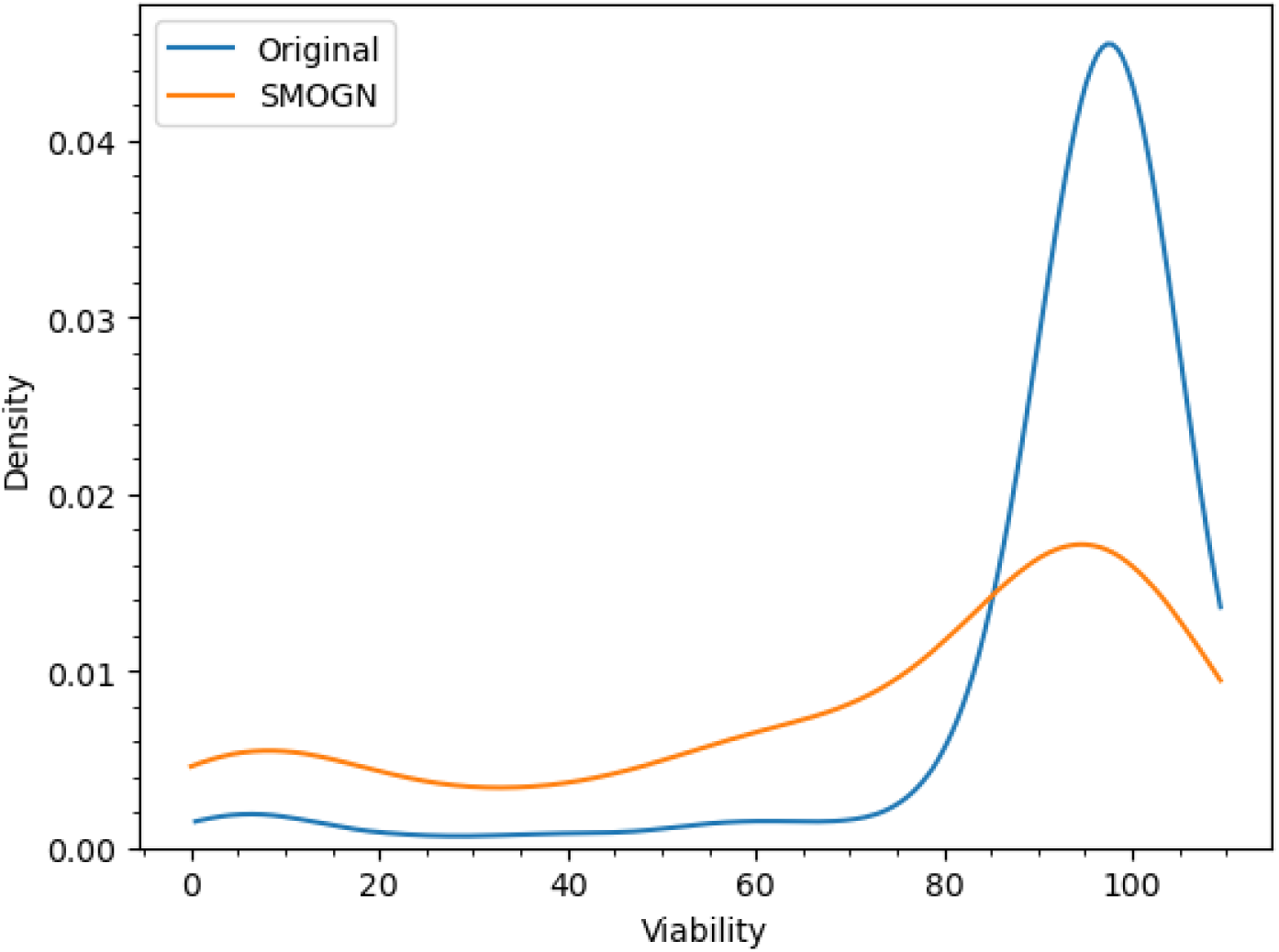
Density plot of the distribution of viability values for C2C12 myoblasts treatment-level data, before data augmentation (blue, Original) and after data augmentation using the SMOGN algorithm (orange, SMOGN).

